# The impact of normal tissue density on tumor growth and evolution in a 3D whole-tumor model of lung cancer

**DOI:** 10.1101/2025.09.25.676637

**Authors:** Rafael R Bravo, Mark Robertson-Tessi, Scott Antonia, Jhanelle Gray, Amer Beg, Robert Gatenby, Matthew B. Schabath, Alexander R A Anderson

## Abstract

Serial low-dose computed tomography (LDCT) scans in patients who are diagnosed with lung cancer during screening offer a history of the densities of tumors and the tissues that surround them during carcinogenesis and cancer progression. We built a CT-scan-resolution computational model to explore how variations in lung tissue density impact tumor growth and evolution in non-small cell lung cancer (NSCLC). Our findings indicate that tumors spread more rapidly through denser tissues when they upregulate glycolysis whilst tumors spread more rapidly through sparser tissues when they upregulate angiogenesis. We used data and images from the National Lung Screening Trial to calibrate our model for untreated lung cancer growth in patients and observed consistency with model predictions in low-density environments.

**Significance:** Our lung lesion model supports prior studies that find tumors tend to evolve toward angiogenic or glycolytic phenotypes. We demonstrate that these evolutionary strategies may be driven by the surrounding normal tissue density and may be observable on imaging.

## 1 Introduction

Medical imaging (e.g., computed tomography [CT]) is standard-of-care in oncology for early detection, diagnosis, treatment planning, monitoring, and image-guided interventions. CT scans are non-invasive x-ray scans that provide 3D volumetric visualization of tissue density at mm-scale resolution. Serial CT scans are imperative for treatment decision-making, as they track whether nodules are growing or shrinking and can detect new metastases. The primary limitation of CT scans is that only tissue-scale density is detected, so determining the cellular or phenotypic composition of a tissue is not directly possible. Furthermore, detecting low-density tumor margins on CT can be difficult.

However, CT scans do capture tissue density heterogeneity in time and space, and we propose that this heterogeneity likely causes different environmental pressures on tumor growth [1–5]. There is evidence that breast cancer cells are responsive to surrounding tissue density; requiring more energy to move and upregulating glycolysis in response to being trapped by dense tissue [6, 7]. To investigate this dynamic in the context of lung cancer, we constructed and calibrated a spatiotemporal population-based model of tumor expansion and evolution within surrounding lung tissue.

We hypothesize that the density of the tissue a tumor invades will select for specific tumor cell phenotypes that can best exploit it; low-density homeostatic tissues will select for different tumor cell phenotypes than high-density tissues. The ultimate goal of this work is to improve patient-specific therapy by inferring the tumor phenotype composition and surrounding tissue context based on the available imaging data. This would predict treatment response dynamics [8–10], and could be used to extrapolate tumor spread beyond detection limits to improve surgical margins and biopsy regions of interest [11].

We used serial CT images from the National Lung Screening Trial (NLST) [12] to calibrate the model and assess the viability of our predictions at capturing the development of real tumors in their native tissue context. The NLST provides a rare glimpse into the initially untreated development of lung cancer in the early detection setting, making it an ideal resource for testing our tumor initiation model. We utilized a subset of data from participants in the annually screened Low-Dose CT (LDCT) arm of the NLST who were diagnosed with NSCLC to model pre-treatment tumor growth dynamics.

## 2 Results

We first summarize our modeling approach and assumptions, then present a series of *in silico* experiments that examine how our model responds to wound healing in low and high-density tissue, how a tumor would grow in such a tissue, and how it might evolve metabolically in different heterogeneous environments. We then characterize patient-specific model calibrations to 3D CT data from 30 NLST patients who developed cancer, revealing the heterogeneity in phenotypes predicted by the model.

### 2.1 Model Overview

Our model simulates the interaction between the tumor and the normal tissue that surrounds it. Each voxel in our 3D spatial domain represents a cubic millimeter of tissue, which is composed of populations of tumor and normal cell types that interact by competing for space and glucose. To initialize the model, we start by using the densities from a normal lung CT scan to set the density of the normal tissue, which is made of epithelial cells (resident tissue) and endothelial cells (vasculature) (Figure 1A). We then seed a tumor that has the capacity to alter its environment using two well-documented phenomena: increased glucose consumption (glycolysis) and tumor angiogenesis [13]. These traits allow the tumor phenotypes to affect the epithelium via competition for glucose and the endothelium through angiogenesis (Figure 1B). Through phenotypic evolution, the tumor can shift the balance of these traits and modify production and consumption of glucose to its potential benefit. Our choice of these phenotypes builds upon our previous work examining both tumor metabolism [4, 14, 15] and angiogenesis [4, 16].

**Figure 1.**
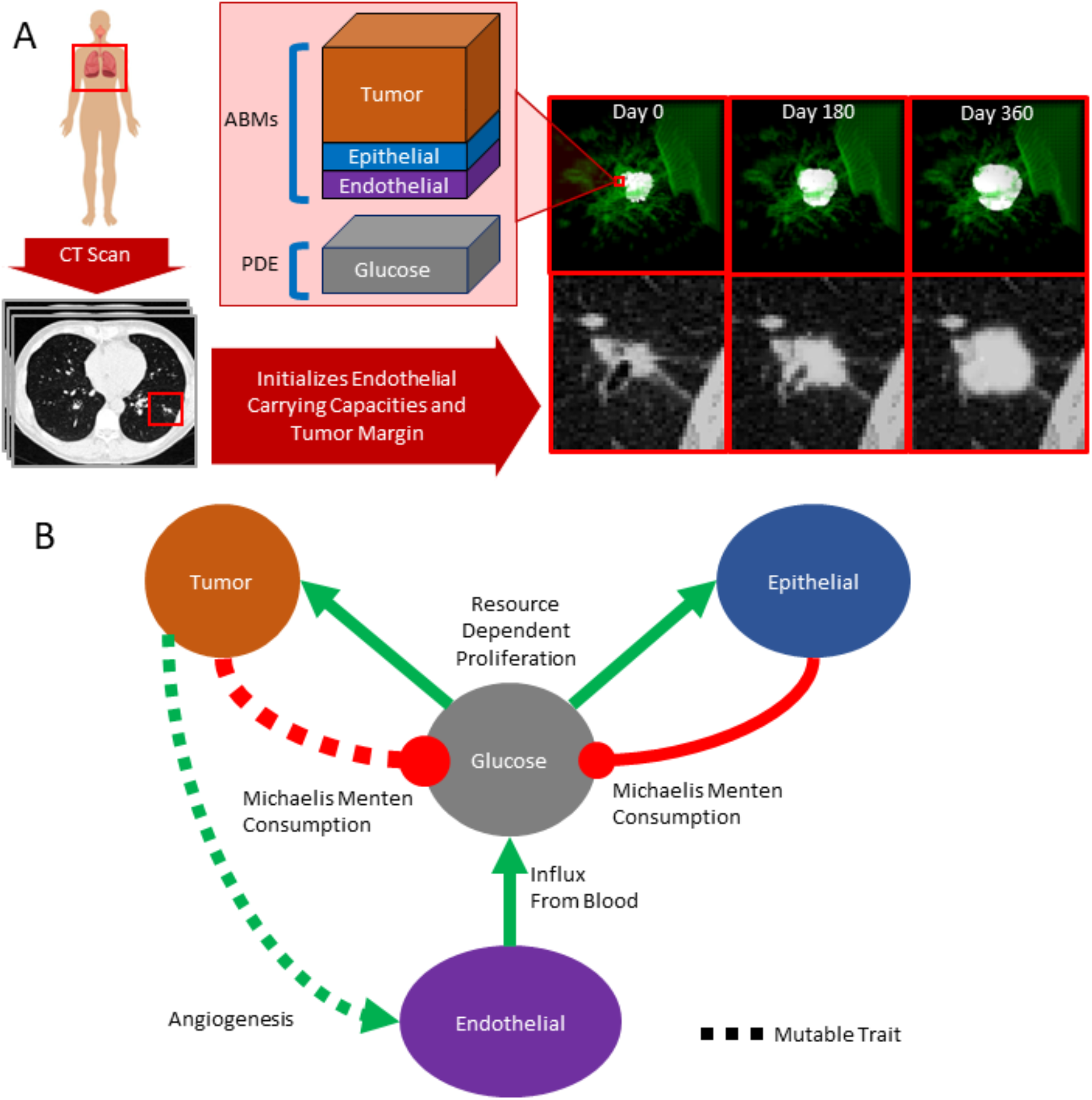
Using CT data to initialize our 3D whole-tumor model of lung cancer. **A** A volumetric CT scan taken from a patient is used to initialize the endothelial cell carrying capacities of each voxel of our model. This endothelial compartment provides glucose for the normal tissue, generating a steady state normal tissue population proportional to the density of the original CT scan. A tumor is then seeded in the center of this environment and grows. We also include the same time points but viewed in 3D, the margin of the high-density normal tissue (500,000 cells or more) are colored in transparent green, while voxels with one more tumor cell are colored white. Below this view, are “virtual CTs” showing the cell density of a central 2D slice through the simulated tumor. Each voxel in the model is composed of discrete counts of cells and a glucose concentration. **B** The interaction network for the components of our model in each voxel, green (red) lines indicate promoting/production (inhibitory/consumption) interactions. Endothelial cells (vessels) deliver glucose, which is consumed by both tumor and epithelial cells, and impacts proliferation rates. Tumor cells utilize angiogenesis to recruit more blood vessels. Dashed lines indicate how the tumor phenotype can change due to mutation.

### 2.2 Wound healing experiment

One of our early goals in building this model was to simulate a homeostatic heterogeneous tissue into which a tumor could grow and evolve. To test the normal tissue behavior in the model, we conducted a simplified wound healing experiment. To initialize the normal tissue in our model, we first took an example patient from the NLST dataset [12] and chose a 3x3x3cm cubic region of normal tissue as the model domain (Figure 2A,B). We visualized cell and diffusible densities at three time points, shown as vertical columns. We created a 0.5cm diameter hole (removed all cells) in the center of this domain, and then observed how the tissue regrew (Figure 2C). After 600 days, the tissue had nearly returned to homeostasis. The overall cell density, labeled “virtual CT” is displayed in the top row, followed by the density of glucose and the density of individual cell types in the subsequent rows.

**Figure 2.**
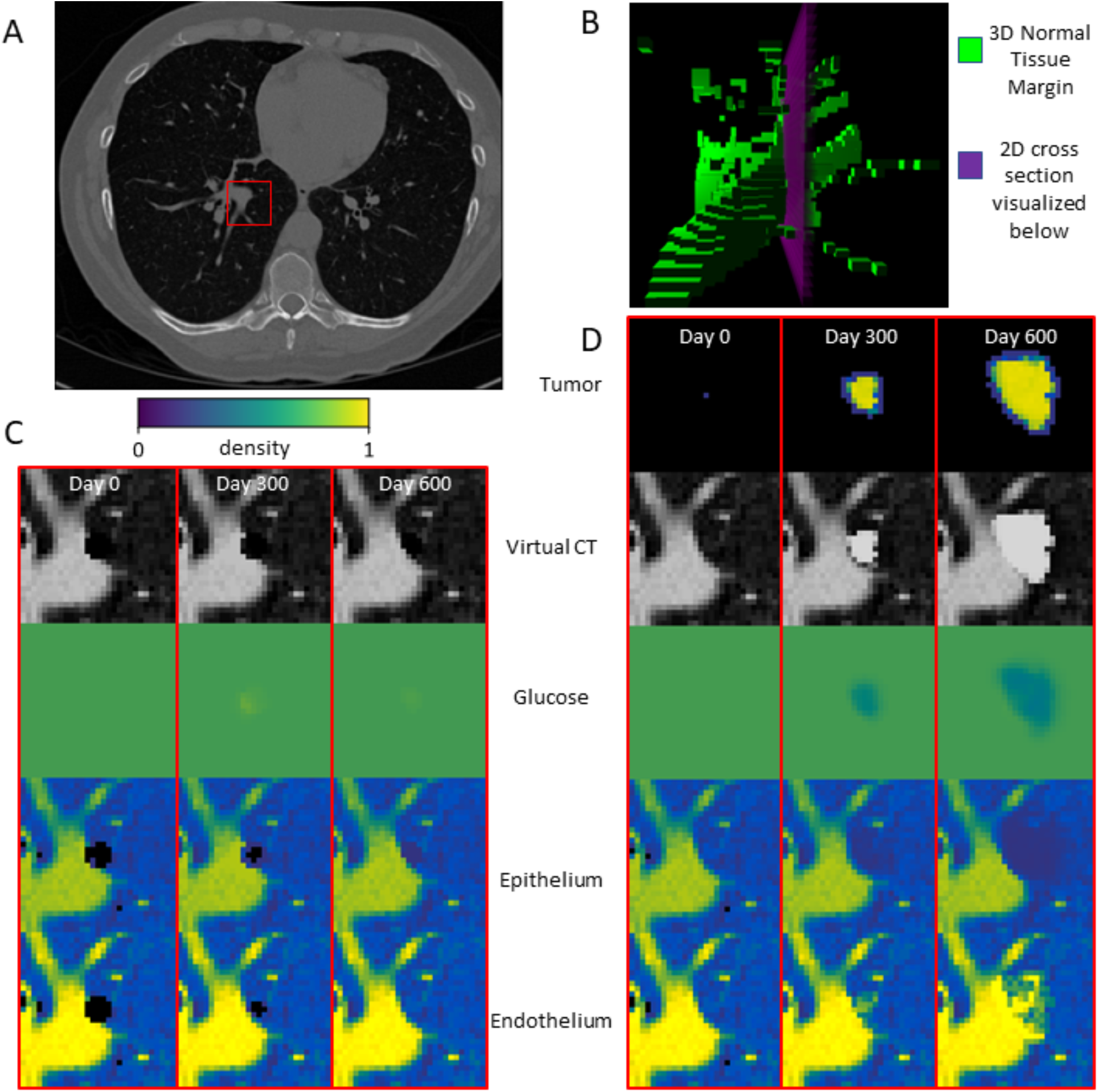
**A** A CT scan from the NLST dataset. The 3D region chosen for this example is indicated with a red square. **B** 3D visualization of the simulation domain. The high-density normal tissue is shown in green. A rectangular 2D slice from the domain (shown in purple) is taken from the 3D model to generate the 2D plots in C and D. **C** A 0.5cm diameter hole is created in the center of the model domain. This hole is refilled by endothelial and epithelial cells over time. Rows show visualizations of the different model components. **D** Example of tumor growth starting with 1000 cells at the center voxel on day zero. The tumor phenotype is (1,1).

We next introduced a tumor to demonstrate the interaction between the tumor and host tissue. We initialized the tumor with 1000 cells in the central voxel, assuming that the tumor cells all shared the same fixed (1,1) phenotype. The first digit indicates activated angiogenesis and the second digit indicates an elevated glucose consumption rate, 2x that of epithelial cells. Tumor cells with a 0 first digit and epithelial cells are not angiogenic. Each further increment of either digit doubles the corresponding phenotypic effect. The impact of this (1,1) phenotype can be seen in the temporal dynamics of the glucose, epithelium, and endothelium in the area of the tumor (Figure 2D). The tumor cell density is displayed in the top row, followed by the same rows as in Figure 2C.

### 2.3 Homogeneous tumor experiments

We next systematically investigated how different tumor phenotypes would be affected by the density of the normal tissue that surrounds them. We started by comparing the growth rates of tumors with different phenotypes across different densities of normal tissue. Our test environment was a 3x3x3cm cubic field of normal tissue at a specified uniform density. For each tumor phenotype, we initially seeded 1000 tumor cells into the central voxel of this domain and observed how long it took for 13,500 voxels (approximately half of the total voxels) to contain at least one tumor cell, which corresponds to when the tumor reaches approximately 3cm in diameter. We measured the growth time for all combinations of our two phenotypic traits (glucose consumption and angiogenic rate) under 6 normal tissue densities (Figure 3A). The angiogenic phenotypes show a growth rate advantage over the glycolytic phenotypes in low density tissue, which inverts in high density tissue.

**Figure 3.**
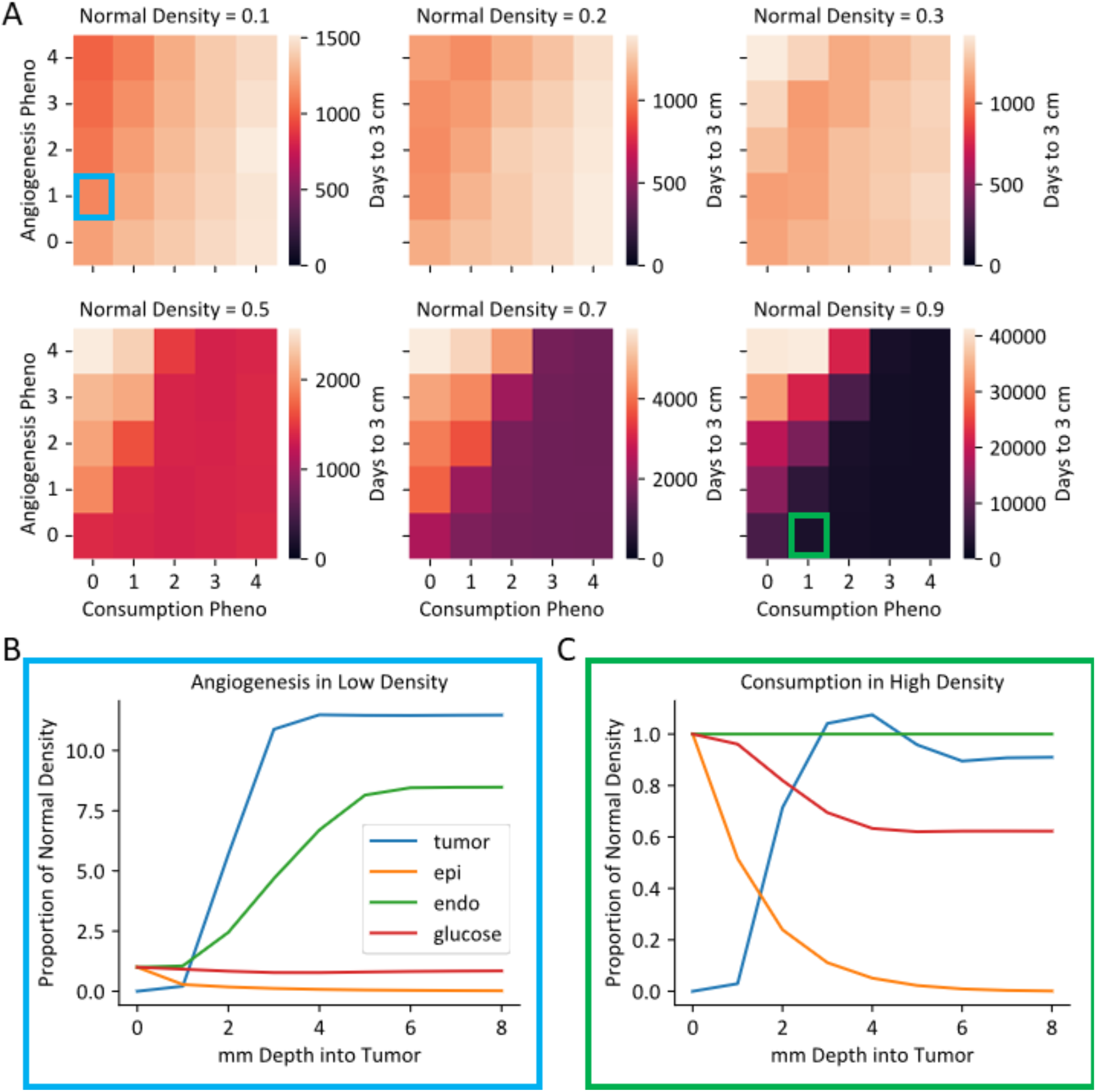
**A** Time for all phenotype combinations to spread into a volume of 13500mm^3^ given different normal tissue densities. **B**,**C** View of average densities of different cell types and glucose at different 1mm-thick layers in the tumor, starting from the tumor edge. Values are normalized by the normal tissue homeostatic densities except for the tumor population which is normalized to the homeostatic epithelial density. The blue and green borders indicate which case from A is being visualized.

We did a more thorough analysis of a select pair of extreme cases by plotting the average densities of the model variables at 1mm intervals (layers) going into the tumor as a proportion of their density in the homeostatic normal tissue. The plotted tumor density is scaled according to the density of the surrounding normal epithelial tissue. Figure 3B shows an angiogenic (1,0) phenotype in a low-density tissue (0.1), figure 3C shows a glycolytic (0,1) phenotype in a high-density density tissue (0.9).

### 2.4 Tumor evolution experiments

We next tested whether the observed growth rate discrepancies between phenotypes in different tissue densities would influence tumor evolution. We gave tumor cells the capability to randomly change phenotype after division in a “mutation event”, which comprises any cell-intrinsic factors that cause a shift in the glucose consumption or angiogenic phenotypes. We tested tumor evolution with a 3x3x3cm field of normal tissue at a uniform density. We seeded 1000 cells into the center of the field with normal tissue glucose consumption and angiogenesis phenotypes (0,0) and observed the tumor phenotypes after the tumor grew to approximately 3cm in diameter. Figure 4A shows 2D slices of tumors evolving at a rate of 0.01 mutations per cell division, revealing substantial spatial heterogeneity.

**Figure 4.**
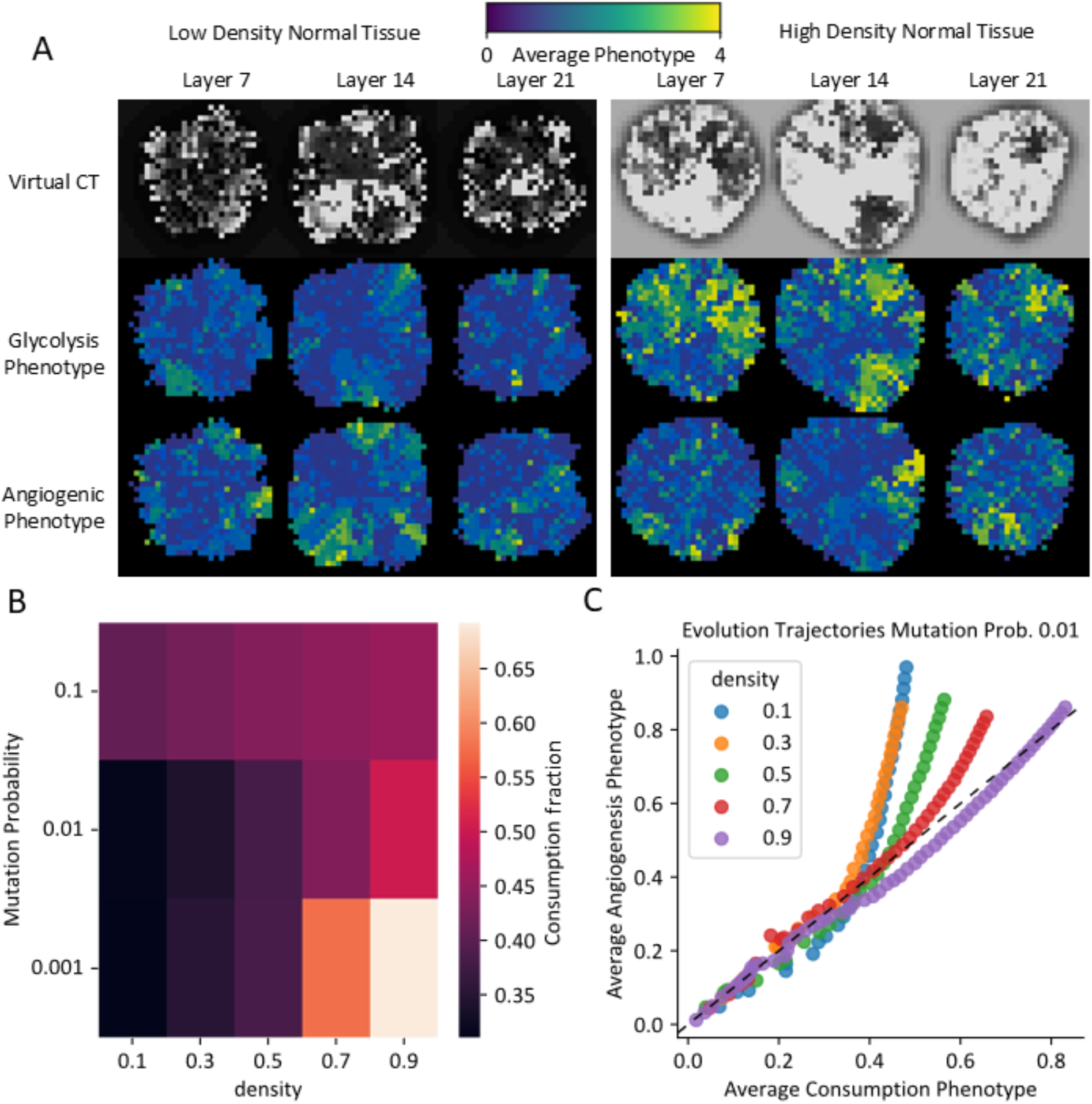
**A** Cross slices of the final tumor states from the tumor evolution experiment. Tumors grew and phe-notypically evolved into different constant densities of normal tissue (left: low-density, right: high-density), producing observably different emergent phenotypes. **B** Fraction of the average glucose consumption phe-notype over total of both average phenotypes indicates how average phenotypes change with density and mutation rate. **C** Average tumor mutational trajectories from the initial (0,0) phenotype under different normal tissue densities. The diagonal line shows the result of random drift.

To better quantify this evolutionary process, we examined what fraction of the tumor was evolving towards a glucose consumption phenotype as opposed to an angiogenesis one. We calculated this fraction by dividing the average glucose consumption phenotype, for all tumor cells, over the sum of the glucose consumption and angiogenic phenotypes. Figure 4B shows a heatmap of these fractions given different mutation rates. We also plotted the average phenotype every 20 days to visualize the average evolutionary trajectory of the tumor (Figure 4C). Average phenotypes start in the bottom left corner at (0,0) and move up and to the right over time. The evolutionary trajectory depends on the tissue density, with the angiogenic phenotype favored in low-density tissue, and the glucose consumption phenotype somewhat favored in high-density tissue.

We next examined how the evolutionary trends found under selection in homogeneous normal tissue density play out in a more realistic heterogeneous lung environment. Taking a 3x3x3cm region of a patient CT scan from the Lung Image Database Consortium [17], we initialized a tumor in two locations: 1) in a low-density lung peripheral region, and 2) in a high-density central bronchial region. We let these tumors grow from an initial population of 1000 baseline phenotype (0,0) cells in a single voxel to a volume of 13500 occupied voxels. The swarm plots in Figure 5A show the average fraction of glucose consumption phenotype that emerged from 40 runs in each starting location. By the end of the experiment the tumors have partially invaded a heterogeneous mix of normal tissue densities. Nonetheless, the density of the starting location has a significant impact on the final phenotype distribution of the tumor.

**Figure 5.**
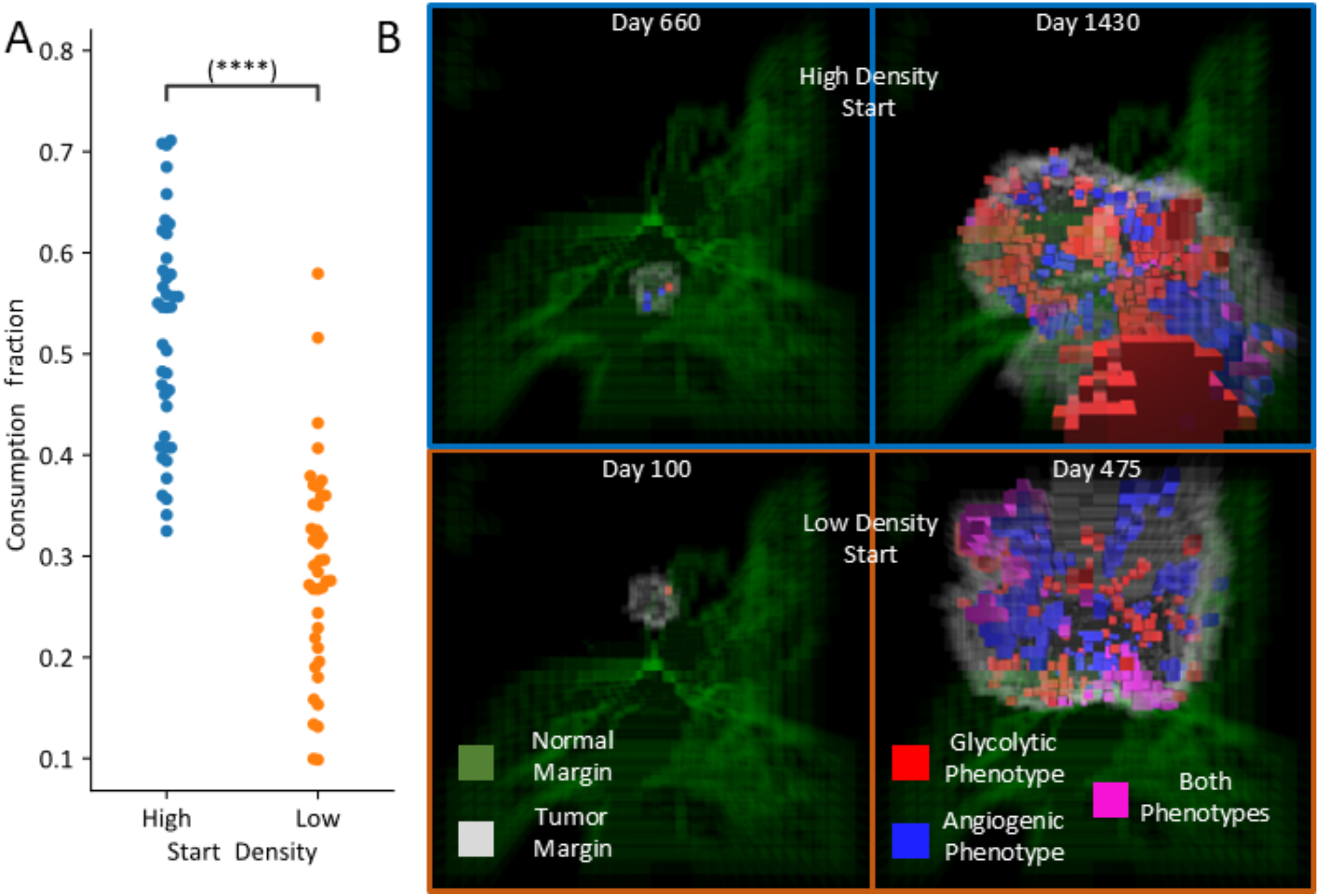
**A** Tumors initiated in different regions of a Lung CT scan. Tumors initiated in high-density regions yielded phenotypes that expressed the glucose consumption phenotype over the angiogenic phenotype, even after breaking out into low-density tissue. Each group contains 40 runs. Statistical significance determined via Mann-Whitney U test. **B** Example 3D visualizations of the tumor invading into a heterogeneous lung tissue from different starting locations. The “bottom” of the 3D domain is closer to the central part of the lung lobe and contains higher density tissue. The margin of normal tissue with density greater than 0.5 is visualized with transparent green voxels. The margin of the voxels containing one or more tumor cells is visualized as transparent white voxels. Average voxel phenotypes whose trait values reach 1 or more are colored based on the average traits that crossed this threshold.

### 2.5 Untreated NLST patient data comparison

In analyzing the NLST dataset, we first approximated the normal tissue density surrounding the tumor by averaging the density of all normal tissue voxels within 3mm of the tumor boundary (Figure 6A). We found that the densities were all lower than 0.3 as scaled by our model. This means that the tumors grew predominantly into low-density tissue.

**Figure 6.**
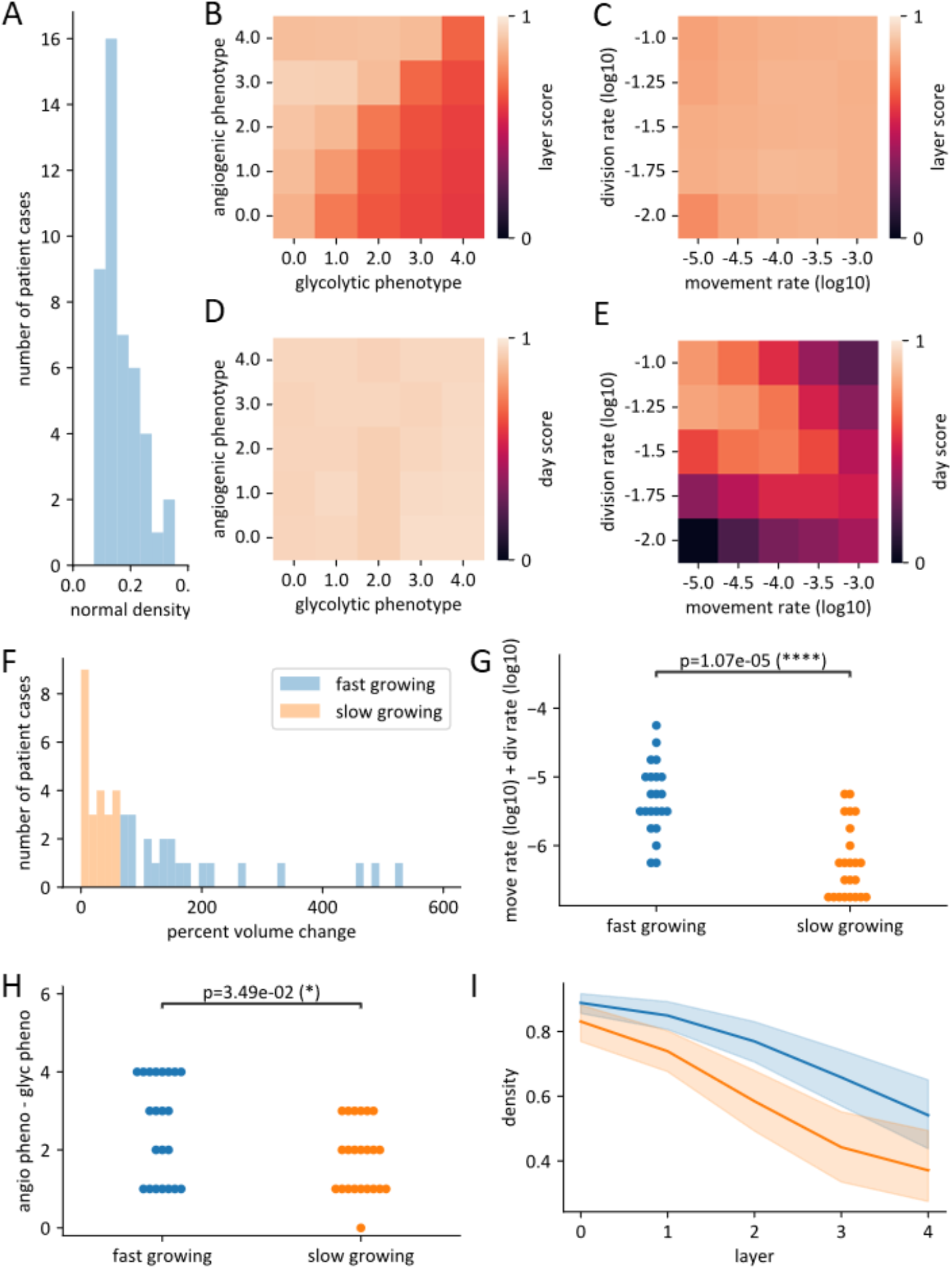
**A** Histogram of average densities of surrounding normal tissue in voxels 3mm or less from the NLST nodules. **B**,**D** Heatmap axes show fixed angiogenic and glucose consumption phenotypes. For each angiogenic-glucose consumption combination, movement and division rates were optimized independently for each patient, and the resulting scores were averaged across all patients. **C**,**E** Heatmap axes show fixed movement and division rates. For each movement-division rate combination, angiogenic and glucose consumption phenotypes were optimized independently for each patient, and the resulting scores were averaged across all patients. **F** Histogram shows patient tumors split into two groups, fast growing (22 cases) and slow growing (23 cases), based on whether growth at 1 year exceeded 65%. **G** The sum of the best fit movement and division rates across the two growth groups. Statistical significance computed via Mann-Whitney U test. **H** Difference between the best glucose consumption and angiogenesis phenotypes for the two growth groups. Statistical significance computed via Mann-Whitney U test. **I** Average tumor density of the first 4 layers of all NLST tumors from the center of the tumor outward for the two growth groups. 95% confidence intervals are shaded.

We next tested whether our model could reproduce several aspects of real tumor growth. We took cases where patients had images of the same tumor that were one year apart, initialized the model with the first image, and ran a parameter sweep of the angiogenesis and glucose consumption phenotypes, the movement probability, and the division probability to find patient-specific tumor parameters that matched the change in tumor density distribution over that year. We thus modeled 45 one-year tumor growth cases from our dataset. From the results of this 4D parameter sweep, we made 2D heat maps (Figure 6B-E) by fixing pairs of the parameters at specific values (shown on the x and y axes), then taking the best scores of the 45 cases given these fixed parameters and averaging them.

Models were scored based on two factors: how close to one year it took for the tumor to grow to the volume of the second CT scan i.e. the day score (Equation 5), and how well the average density of 1mm layers of the tumor matched the average density of the same layers from the second CT scan i.e. the layer score (Equation 6). This allowed us to investigate which parameter values were indispensable for achieving good scores and which could be compensated for by the other parameters. The resulting heatmaps show that the layer score is most sensitive to the angiogenesis and glucose consumption phenotypes (Figure 6B), while division and movement rates do not impact the best score significantly (Figure 6C). In contrast, the day score is insensitive to angiogenesis and glucose consumption phenotypes (Figure 6D), but requires an appropriate balance of division and movement rates to score well (Figure 6E). This indicates that both scoring functions in combination help constrain relevant combinations of tumor growth parameters and phenotypes, with the day score primarily constraining growth and migration parameters, and the layer score primarily constraining angiogenic and glucose consumption phenotypes.

To look in more detail at differences in tumor growth in the dataset, we separated the patient data evenly into two groups: a fast growing group where the tumor volume increased by more than 65%, and a slow growing group where the tumor volume increased by less than 65%. We then looked at which combinations of movement rate and proliferation rate best fit these patients (Figure 6F), and see that faster growing tumors tend to both move and divide faster (Figure 6G). We also see that the faster growing tumors are more angiogenic and less glycolytic (Figure 6H). We further looked at the average layer densities from the center of the patient tumors (most tumors have at least 4 layers) and find that fast growing tumors tend to be denser (Figure 6I), which could explain the difference in best-fitting parameters as in our model denser tumors require increased angiogenesis and decreased glycolysis.

## 3 Discussion

We have simulated tumors at a clinically detectable spatial scale while preserving the ability to track the stochastic behaviors of individual cells. Using our Population-Based Model (PBM) approach we directly simulated realistic numbers of cells in an organ-scale 3D tumor model, allowing us to bridge cell-scale phenotypic changes and interactions up to tissue-scale dynamics with only desktop computing power.

Across all simulations, we find an eco-evolutionary dynamic in which the dominant tumor phenotypes are selected by properties of the normal tissue in which the tumor evolves. Low-density normal tissue selects for an angiogenic phenotype, while high-density normal tissue selects for a glycolytic phenotype. Figures 3B,C demonstrate the reasons for this: in low-density normal tissue, the tumor is bounded by limited access to resources, and through angiogenesis is able to acquire more resources to accelerate growth. In high-density normal tissue, the tumor is bounded by spatial competition with the epithelium, and through increased glycolysis is able to starve the epithelium and decrease the tissue density on the periphery of the tumor, facilitating faster invasion. It is worth noting that in our model tissue density encompasses many co-varying properties that we do not explicitly separate (composition of endothelium and epithelium, for example), which is a simplification of the heterogeneity of normal tissue.

### 3.1 Analogous Tumor Phenotypes

This tendency for angiogenic phenotypes to evolve in low density tissue and glycolytic phenotypes to evolve in high density tissue parallels the differences between squamous cell carcinoma (SCC) and adenocarcinoma (ADC) cell types in NSCLC. ADC tends to emerge in peripheral locations of the lungs [18]. The peripheral regions are composed of the alveoli and capillaries, with large regions of air and therefore a decreased normal tissue density. Interestingly, the antiangiogenic drug bevacizumab when combined with carboplatin and paclitaxel increased median survival in ADC in a phase II clinical trial; however, this benefit was not observed when SCC was included in the cohort [19].

In contrast, SCC tends to emerge in central locations of the lungs [18]. These locations house the bronchi and are made up of dense epithelium, basement membrane, large blood vessels and connective tissue, all of which contribute to higher density. SCC cells tend to be more glycolytic as they upregulate GLUT1 [20, 21] and have been shown to be more sensitive to glucose deprivation than ADC [21]. Although these observations about ADC and SCC are phenomenological and not causal, they are consistent with our results.

Another analogous phenotypic distinction can be found in the EGFR and KRAS subsets of ADC. The EGFR-mutant phenotype in ADC occurs more in younger, non-smoking patients [22] and tends to be more angiogenic than KRAS mutants [23], which tend to be more glycolytic [24]. The KRAS mutant phenotype occurs more frequently in older patients with a history of smoking [22, 25]. These patients also tend to have fibrotic lung tissue, which increases tissue density to the point that it is appreciable in CT scans, and also increased bronchiolar artery vessel dilation and neovascularity [26]. This observation is consistent with our model prediction that higher density tissue favors glycolytic tumor phenotypes.

### 3.2 Evolved Tumor Heterogeneity

From our evolution experiments, we see that the evolution of the tumor phenotype diverges depending on the density of the normal tissue around the tumor, with sparser tissue selecting for angiogenesis expression, and denser tissue selecting for elevated glucose consumption. These trends are sensitive to mutation rate since faster mutation rates cause any context dependent phenotypic selection to be overshadowed by drift (Figure 4B).

We also see spatial phenotypic heterogeneity readily develop in our evolutionary model. This may be because these phenotypes primarily modify invasion into normal tissue, but have a lesser impact on direct competition between phenotypes, allowing random drift to strongly influence tumor composition. It has previously been observed that while one phenotype is dominant during the early stages of cancer development, they co-exist in larger tumors. That is, the glucose consumption phenotype is dominant in the invasive edge of the tumor while the angiogenic phenotype is dominant in the tumor core[15, 27]. Furthermore, therapeutic perturbations could alter these eco-evolutionary dynamics to promote, for example, proliferation of the angiogenic phenotype at the expense of the glycolytic population [28].

We also saw that the tissue of origin impacts the average tumor phenotype even after it encounters other contexts (Figure 5). We observed that when starting in high-density tissue, the tumor develops a more glycolytic phenotype to facilitate invasion through the dense tissue context. Once the tumor escapes, it more quickly invades the low-density tissue, leaving less time and pressure for shifting the phenotype back towards angiogenesis. When starting in low-density tissue however, angiogenesis is selected for from the beginning, and invasion into high-density tissue happens so slowly by comparison that the angiogenic phenotype dominates.

### 3.3 NLST Fitting Implications

In our untreated NLST patient-data comparisons, we find that proliferation and migration parameters are the most important factors for fitting the observed tumor growth rates, whereas the angiogenic and glucose consumption phenotypes are more important for fitting the tumor layer density distributions seen on imaging (Figure 6B-E). This suggests that the layer density distribution of a patient tumor may be a potential biomarker for its angiogenic or glucose consumption state. We find that tumors with greater angiogenic expression and less glucose consumption tend to match the faster growing patient cases, which matches our expectation for tumors growing into a lower-density environment (Figure 6C).

While we did find this concordance with our model in a low-density tissue context, we did not see examples of tumors growing into high-density tissue in our dataset and were thus unable to contrast both tissue environments with NLST examples. There is evidence that it is more difficult to detect tumors growing into central lung tissue with CT [29], which could partially explain this, although other aspects of early NSCLC development or cohort composition may also contribute.

Whilst our focus here was on lung cancer invasion, our PBM approach is more broadly applicable. Temporal CT dynamics are a potentially rich clinical data source that could be exploited to better understand the impact of treatment as the therapy happens. Organ-scale models directly connect with this scale of data and our PBM approach allows us to connect with cell-level heterogeneity observed at the histological scale from imaging guided biopsies [30], for example. The ability to integrate clinical data from different spatial and temporal scales is going to be an important part of future clinical decision-making and one we believe our PBM approach will play an important role in.

## 4 Methods

### 4.1 Population-based model

To match the imaging data directly, the model must accommodate tumors large enough to be detected with CT imaging, ranging from a few millimeters to several centimeters in radius. A cellular ABM at this scale must simulate all the cells in such a macroscopic tumor (at least 100 million cells), which is computationally prohibitive. Therefore, we developed a novel PBM that tracks localized and discrete cell populations, at the voxel scale, rather than individual agents at the cell scale. Each population in the PBM has a distinct state defined by cell count, phenotype, and position. PBM dynamics are based on probabilistic rules that alter cell counts by adding or subtracting cells, or moving cells between voxels and/or phenotypes. These probabilities are resolved into counts stochastically using an efficient binomial and multinomial distribution sampler that is incorporated into HAL [31]. Consolidating agents into populations also makes the computational cost of simulating a PBM dependent upon the number of distinct populations, rather than the number of agents. Thus, we can simulate an arbitrary number of agents per voxel, and have chosen a maximum of 1 million cells per 1mm^3^ as an appropriate maximum density per voxel [32].

A more common approach to building organ-scale tissue models is to use partial differential equations (PDEs) to represent the density of cells over space and time [3, 33]. A major difference with PDEs is that they are continuous and generally deterministic, whereas PBMs are discrete and stochastic. This method was initially developed as part of our Hybrid Automata Library (HAL)[31], a software library for hybrid agent-based modeling, and has been independently developed in other publications. [34, 35]

### 4.2 Model components

Each voxel in the model contains a glucose concentration and counts of several different cell types, which are summarized in Figure 1B. The normal cell types that comprise the model are endothelial cells and epithelial cells. These cell populations begin at a glucose-driven equilibrium, with endothelial cells producing glucose and the epithelial cells consuming glucose such that the average steady-state concentration of glucose in the tissue is around half that in the blood [36]. To generate homeostatic normal tissue at a specific density, we assume that there is a required vascular density in normal tissue to support a given tissue density. This leads to a ratio of 1 endothelial cell for every 33 epithelial cells [37]. This naturally limits the endothelial population and therefore the amount of glucose influx from the blood. In the absence of tumor induced angiogenesis, the epithelial population maintains a homeostatic tissue dictated by glucose availability. The baseline endothelial carrying capacity *c*_*i*_ at each position *i* is scaled linearly between 0 and the maximum baseline endothelial carrying capacity *c*_*b*_, depending on the desired tissue density.

The tumor cells behave similarly to the epithelial cells. However they will divide even when glucose is below normal levels, leading to abnormal growth and competition with the epithelial cells. The tumor cells may also mutate and become angiogenic to recruit more endothelial cells, and/or become more glycolytic, consuming glucose faster. This is an over-simplification of the metabolic processes that help regulate normal tissue homeostasis, however, we have deliberately chosen to not include complex metabolic processes including the role of other factors such as oxygen or acidity to reduce model complexity.

### 4.3 Day loop overview

The model iterates with a step function that simulates one day per iteration. The function has 5 phases summarized in Figure 7 and Figure S1. (i) cell death; (ii) cell division; (iii) tumor cell mutation; (iv) cell movement; and (v) glucose dynamics in response to these changes. We next discuss the phases in detail and the equations that govern them. The parameters for these equations are summarized in Table S1.

**Figure 7.**
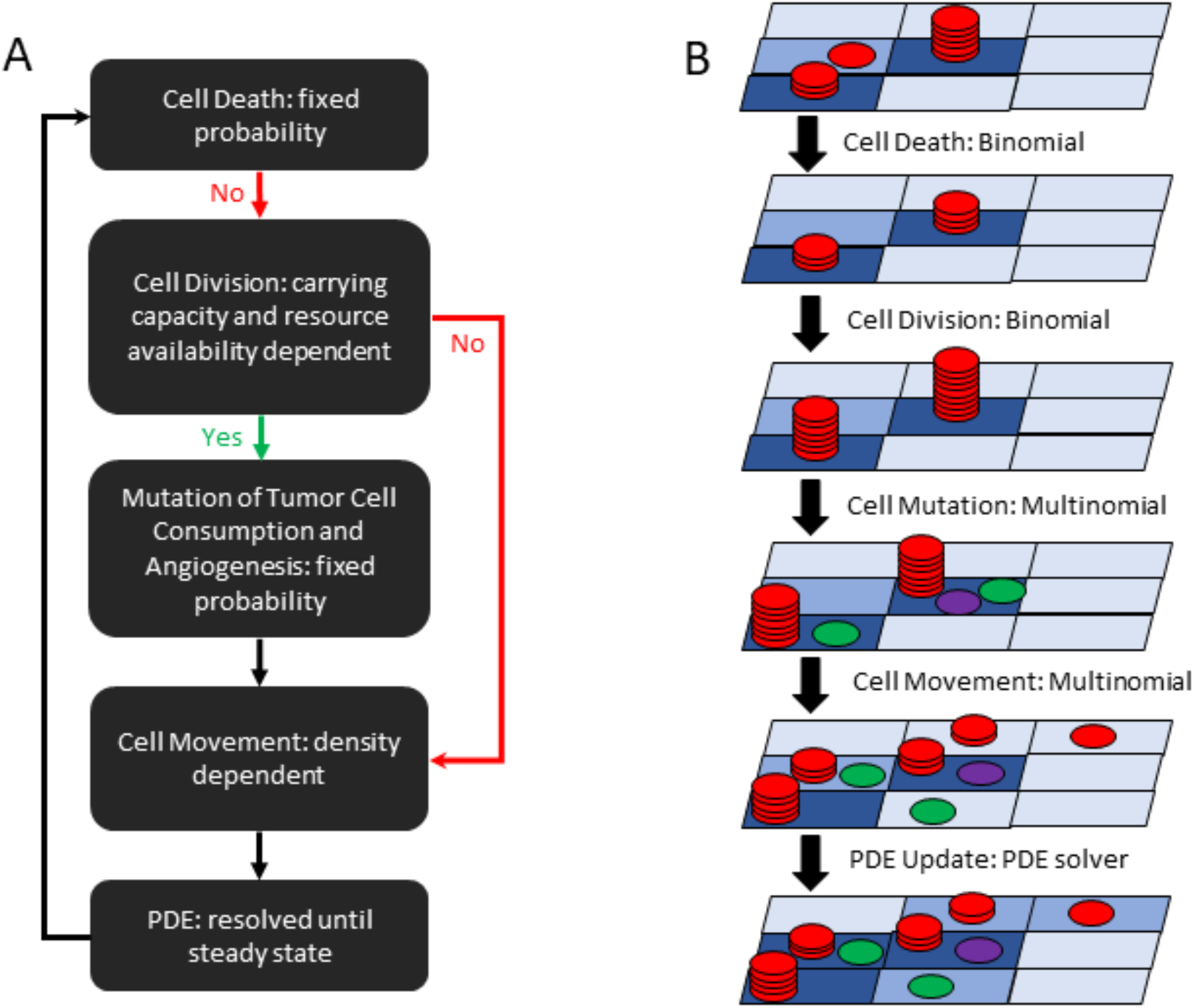
**A** Diagram of the Step function day cycle. This cycle is repeated for every simulated day. Updates to cell counts are made to the full domain simultaneously using the binomial and multinomial. **B** Image representation of the step cycle. Cells are represented as circles, while PDE concentrations are represented as differently colored rectangles. The text next to the arrows indicates what updates are applied between cycle states.

### 4.4 Death and division

First, a fixed random death probability rule is applied to all cell populations. This models the natural turnover of tissue [38] and tumor cells. Cells that survive this step have a chance of proliferating.

Proliferation is affected by multiple limiting rules. The rule that yields the lowest probability sets the proliferation probability. The first proliferation rule enforces a voxel carrying capacity to model physical crowding limitations. For a population of cell type *p* at position *i*, this rule (Equation 1) maximizes proliferation when the voxel is unoccupied and decreases the proliferation probability exponentially to zero as the sum of all normal and tumor populations at the voxel *N*_*i*_ approaches the spatial carrying capacity *k* of 1 million cells per voxel. The exponent *p*_*x*_ (2) defines how quickly the probability decreases as the population increases. The maximum proliferation probability per day *p*_*p*_ is cell-type dependent (0.1 for normal cells, 0.25 for tumor cells):

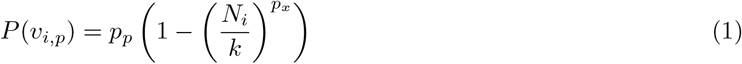

The second proliferation rule (Equation S7) reflects glucose availability for the epithelial and tumor cells. This proliferation probability hits a maximum value for epithelial and tumor cells at 80% and greater blood glucose concentration and drops linearly to zero at 50% blood concentration for epithelial cells. This rule causes the epithelial cells to maintain a homeostatic population with approximately 50% blood glucose concentration in the tissue [36]. The tumor cells will keep proliferating until a minimum of 30% blood concentration of glucose, which along with a greater max proliferation rate are the main growth advantages that the tumor cells have over the epithelial cells (Figure S2).

The endothelial cells follow the same voxel carrying capacity rule, but ignore glucose and have an endothelial-specific carrying capacity rule, which is set proportionally to the tissue density in the original CT scan (Equation S8). This generates the appropriate resource distribution to support the density of normal cells matched to the original CT scan, thus facilitating the emergence of the underlying normal lung tissue structure. The endothelial carrying capacity can also be altered by angiogenesis, which depends on how large and angiogenic tumor populations are at that voxel (Equation S9). This angiogenic population is assumed to proportionally increase the original endothelial carrying capacity at that voxel until an upper bound of 4 times the normal maximum density endothelial capacity is reached [39].

### 4.5 Tumor evolution

Following division, tumor cells can shift their phenotype via random mutation of two traits: Glucose consumption rate *r*_*g*_ (Equation S10) and angiogenesis rate *v*_*a*_ (Equation S11). This is a stochastic event that occurs with probability 0.01 during cell division for each trait independently and can lead to increases or decreases in these traits by one increment. Each trait can shift between 5 states, which we will represent with coordinate pairs (*a,g*). At state (0,0), the glucose consumption and angiogenesis rates are equivalent to that of epithelial cells: a normal level of glucose consumption and no angiogenesis. With the first increment of the angiogenic trait, tumor cells become angiogenic. Each further increment increases the angiogenesis rate *v*_*a*_ by a factor of 2. Each increment of the glucose consumption phenotype increases the consumption rate of glucose *r*_*g*_, by a factor of 2.

### 4.6 Movement

We next describe the cell movement probability function *P* (*m*_*i,j*_), which models the impact of migrating through denser tissues by decreasing the probability as a power of cell density, like the proliferation probability function. Cells are impeded as a function of the average number of cells between the voxel that they occupy *N*_*i*_ and the voxel that they are attempting to move into *N*_*j*_. See Equation 2 where *k* is 1 million cells and *m*_*x*_ is 2.

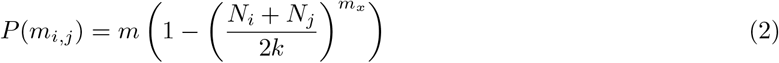

The migration function is applied using the 19-point neighborhood in 3D, allowing migration into the 18 voxels in the Moore neighborhood around the voxel of origin, excluding the 8 corner voxels. The migration rates into each voxel of the neighborhood are set proportional to their distances from the current voxel, capturing the slightly reduced likelihood of movement over the larger distances. This also leads to smoother and more realistic tumor population expansion.

### 4.7 Glucose dynamics

We next describe the PDE (3) governing the glucose component of the model [10] at position *i*. Glucose is produced by endothelial cells *V*_*i*_, diffuses with rate *g*_*d*_, and is consumed by epithelial cells *E*_*i*_ and tumor cells. The glucose production rate *g*_*p*_ is proportional to the number of endothelial cells and decreases linearly as the tissue glucose concentration approaches the blood glucose concentration *g*_*b*_ in accordance with Fick’s law [40]. Glucose is consumed by epithelial cells *E*_*i*_ with rate constant *r*_*e*_ and by tumor cells. The total number of tumor cells in voxel *i* with glycolytic phenotype *g* is denoted *T*_*ig*_. These populations consume glucose with rate constant *r*_*g*_, which depends on glycolytic phenotype. Consumption follows Michaelis–Menten dynamics with half-max constant *g*_*h*_ [41, 42]. The glucose PDE is resolved to steady state at each time step by iterating the PDE update function until the maximum change across all voxels between iterations drops below a tolerable threshold *ϵ*.

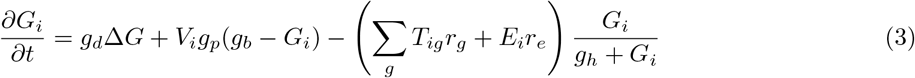

### 4.8 CT scan integration

The tumor margins for this dataset were segmented in a prior study [37] and were used for the work conducted here. Only cases where the patient nodules were segmented over subsequent years and the patient was later diagnosed with either SCC or ADC were considered. Affine alignment of the scans across time points was done prior to our analysis using the SimpleITK package [43] to maintain scale and spatial orientation. After excluding a handful of scans that were misaligned or where the nodule shrunk between years, we had a total of 48 cases where a segmented nodule grew over a 1 year period (Figure S3A,B). To improve the performance efficiency of our analysis, we excluded two tumors with radii 28mm and 30mm (all other nodules had radii from 7mm to 22mm), resulting in 45 cases from 30 patients. Additional analysis of this group can be found in Figure S4. The tissues are initialized such that their spatial volumetric densities match the first scans (Figure S3C), with tumors established within the segmentation margins, and then the tumors are allowed to grow over time until the simulated tumor volumes match or exceed the measured tumor volumes at the second scans (Figure S3D) one year later.

To initialize the model using CT data, we first resolve the CT voxels into 1mm^3^ via interpolation, which we accomplish using the SimpleITK package [43]. We next take a 3x3x3cm cubic region of the interpolated tissue to define the model domain. To input the CT densities to the model, we establish a linear mapping between Hounsfield units (HU) from the CT to a count of cells per mm^3^, with -1000 Hounsfield units (HU) mapping to zero cells and 100 HU mapping to a million cells [44]. We assume the earlier established homeostatic ratio of 1 endothelial cell for every 33 epithelial cells [37]. Thus the local endothelial cell carrying capacity is set proportional to the HU value at a given voxel and maintains the epithelial cell population at the desired density.

A tumor population can be added to this system either at a central voxel as a seed population of cells, or by using a tumor segmentation that accompanies the CT scan. To use a tumor segmentation, we need to divide the segmented voxels into normal and tumor cells. We set the density of normal cells within the tumor region to be the same as the average of the surrounding tissue within 3mm of the segmentation margin, and add tumor cells to make up the rest of the voxel density. If the tumor is angiogenic, the endothelial cell counts within the tumor region are increased to the voxel carrying capacities imposed by local tumor angiogenesis.

Models were scored (Equation 4) based on two factors: how close to one year it took for the tumor to grow to the volume of the second CT scan (Equation 5) i.e. the day score *s*_*y*_, (where *d* is the number of simulated days it took for the model tumor to grow to the volume of the second scan) and how well the average density of 1mm layers of the tumor matched the average density of the same layers from the second CT scan (Equation 6) i.e. the layer score *s*_*d*_, (where *n* is the number of tumor layers compared, *c*_*l*_ is the average density of a given layer of the model tumor, and *m*_*l*_ is the average density of a given layer of the patient data tumor) see Figure S3E for more details. If the number of layers is inconsistent between the model and the data, then *n* is set to whichever layer number is largest and the *c*_*l*_ − *m*_*l*_ term is set to 1 for the mismatched layers to penalize the mismatch.

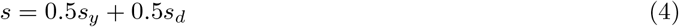

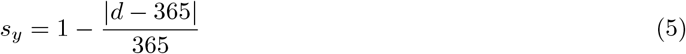

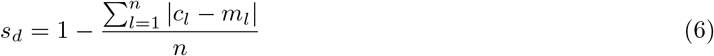

We had to modify some of the model parameters in order for the tumor voxel densities and growth trajectories to match between the model and the patient data. We found that the tumors in the dataset reached a maximum density near 1.0, while the model tumors reach maximum density nearer to 0.8 due to contact inhibition. To adjust for this, the contact inhibition on the cells was reduced by lowering the proliferation exponent *p*_*x*_ from 2 to 1. We also found that the tumors from the dataset tended to grow more slowly than the model tumors. This meant that the fitted tumor proliferation rates were usually significantly lower than our earlier experiments. To accommodate the lower growth rate, such that the tumor could still compete with the epithelial cells, the death probability of the tumor cells was adjusted to be equivalent to the death probability of the epithelial cells. To initialize the normal and tumor tissue, we used the segmentations from the NLST dataset to define the tumor region. See the CT Scan Integration section of the methods for details.

### 4.9 Data Availability

The NLST patient data used in this study are not publicly available due to patient privacy requirements. Other data generated in this study are available within the article and its supplementary data files.

### 4.10 Declaration of generative AI and AI-assisted technologies in the manuscript preparation process

During the preparation of this work, the authors used ChatGPT to suggest edits that improved clarity and readability. The authors reviewed and edited the output as needed and take full responsibility for the content of the published article.

## Notes

Authors have no conflict of interest to declare

https://github.com/MathOnco/LungTissueDensityModel

### Competing Interest Statement

The authors have declared no competing interest.

### Summary of Updates

Updated manuscript to reflect the version submitted for peer review, including revisions to the text, figures, and supplementary materials.

### Funding

- National Cancer Institute via the Cancer Systems Biology Consortium (CSBC): U54CA274507
- Moffitt Center of Excellence for Evolutionary Therapy

