## Supplementary Materials for "The impact of normal tissue density on tumor growth and evolution in a 3D whole-tumor model of lung cancer"

### 1 PBM rules expanded

| Parameter | Symbol | Units | Value | Source |
|---|---|---|---|---|
| Normal Max Proliferative Probability | $p_e, p_v$ | P/day | 0.1 | [45] |
| Tumor Max Proliferative Probability | $p_t$ | P/day | 0.25 | Model-Specific |
| Proliferation Exponent | $p_x$ | - | 2 | Model-Specific |
| Migration Exponent | $m_x$ | - | 2 | Model-Specific |
| Epithelial Death Probability | $d_e$ | P/day | 0.00625 | [38] |
| Endothelial Death Probability | $d_v$ | P/day | 0.000435 | [38] |
| Tumor Death Probability | $d_t$ | P/day | 0.0125 | Model-Specific |
| Spatial Carrying Capacity | $k$ | cells/mm$^3$ | 1E6 | [32] |
| Blood Glucose Concentration | $g_b$ | mol/L | 5.56E-5 | [46] |
| Maximum Proliferation Glucose Threshold | $h_m$ | mol/L | 0.8*5.56E-5 | Model-Specific |
| Epithelial Proliferation Glucose Threshold | $h_e$ | mol/L | 0.5*5.56E-5 | [46] |
| Tumor Proliferation Glucose Threshold | $h_t$ | mol/L | 0.3*5.56E-5 | Model-Specific |
| Maximum Baseline Endothelial Carrying Capacity | $c_b$ | cells/mm$^3$ | 3E4 | [39] |
| Maximum Endothelial Carrying Capacity | $c_m$ | cells/mm$^3$ | 1.3E5 | [39] |
| Cell Migration Rate | $m$ | mm$^2$/day | 1E-6 | Model-Specific |
| Glucose Diffusion Rate | $g_d$ | mm$^2$/sec | 0.002 | [47] |
| Epithelial Glucose Consumption Rate | $r_e$ | mol/sec | 2.5E-19 | [48] |
| Endothelial Glucose Production Rate | $g_p$ | mol/sec | 4.5E-9 | Model-Specific |
| Base Angiogenesis Effect | $v_1$ | - | 0.0303 | Model-Specific |
| Half-Max Glucose Constant | $g_h$ | - | 0.5 | Model-Specific |
| Mutation Probability | $u$ | P/division | 0.01 | Model-Specific |
| Phenotype-Specific Glucose Consumption Rate | $r_g$ | mol/sec | $r_e 2^g$ | Eq. S10 |
| Phenotype-Specific Angiogenesis Effect | $v_a$ | - | Eq. S11 | Model-Specific |
| CT Minimum Density | - | Hounsfield Units | -1000 | [44] |
| CT Maximum Density | - | Hounsfield Units | 100 | [44] |

Supplementary Table 1: Model parameters used throughout experiments unless otherwise specified.

#### 1.1 Glucose dependent proliferation rule

The second proliferation rule for epithelial and tumor cells reflects glucose availability. This proliferation probability $P(v'_{i,p})$ at position $i$ and for cell type $p$ hits a maximum value at 80% blood glucose concentration ($h_m$) and drops linearly to zero at the proliferation glucose threshold ($h_p$), where $h_p = h_e$ for epithelial cells and $h_p = h_t$ for tumor cells. This rule causes the epithelial cells to maintain a homeostatic population with approximately 50% blood glucose concentration in the tissue [36]. The tumor cells will keep proliferating until 30% blood concentration of glucose is reached. This function is shown in Equation S7 and Figure S2, where $G_i$ denotes the glucose concentration at position $i$.

$$P(v'_{i,p}) = \begin{cases} p_p & G_i \geq h_m \\ p_p \dfrac{G_i - h_p}{h_m - h_p} & h_p < G_i < h_m \\ 0 & G_i \leq h_p \end{cases} \tag{S7}$$

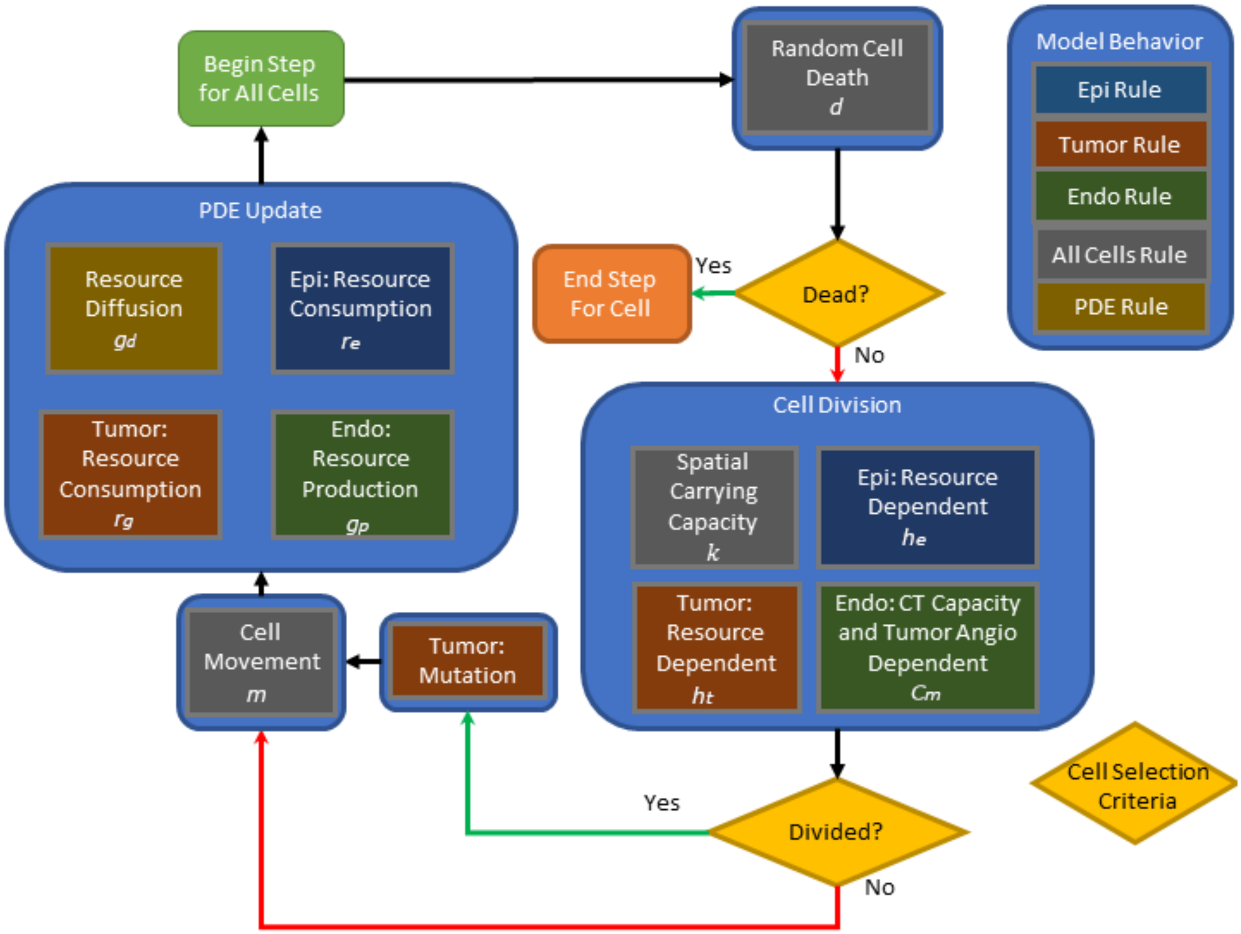


Supplementary Figure 1: Diagram of Step function day cycle logic. This cycle is repeated for every simulated day. Cells undergo random death, those that survive may divide, those that divide may mutate if they are tumor cells, then all surviving cells may move, then the PDEs are run until steady state.

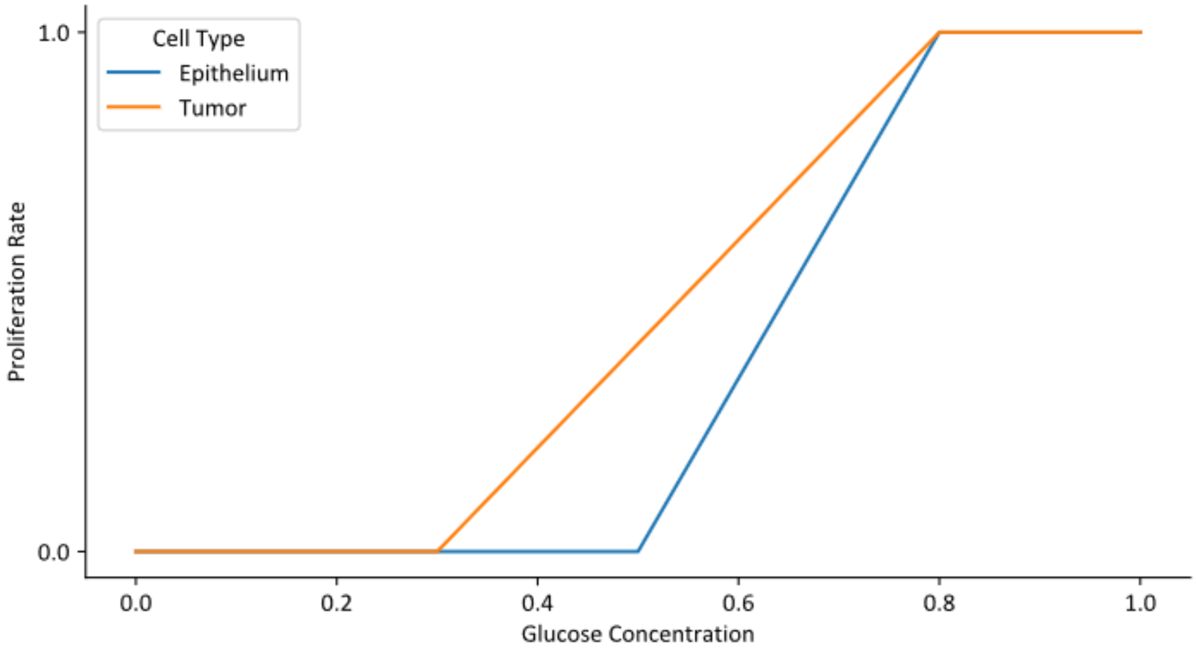


Supplementary Figure 2: Illustrates how glucose concentration and proliferation rate are related for epithelium and tumor. This rule is combined with a contact inhibition effect which can also limit proliferation. The y-coordinates show the proliferation rate normalized by the maximum daily proliferation probability for each cell type: 0.25 for tumor cells and 0.1 for normal cells.

#### 1.2 Endothelial carrying capacity rule and angiogenesis

The endothelial cells don't have a glucose dependent proliferation rule, but have a second carrying capacity rule. The probability of division $P(v_i'')$ of endothelial cells at a given voxel follows a logistic function. The carrying capacity of this function at a given voxel $V_i$ is set proportional to the density at that voxel in the lung CT used for initialization. Equation S8 shows how this endothelial proliferation probability is calculated.

$$P(v_i'') = p_v \left(1 - \frac{V_i}{C_i}\right) \tag{S8}$$

The endothelial carrying capacity $C_i$ at position $i$ can also be altered by angiogenesis from the tumor. Angiogenesis occurs at a rate $v_a$ per tumor cell at position $i$ depending on the angiogenesis phenotype $a$. This angiogenesis factor is added to the original endothelial carrying capacity $c_i$ and allows this capacity to linearly increase until the upper bound $c_m$. See Equation S9, where $T_{ia}$ denotes the total number of tumor cells at position $i$ with angiogenic phenotype $a$.

$$C_i = \begin{cases} c_m & c_i + \sum_a v_a T_{ia} \geq c_m \\ c_i + \sum_a v_a T_{ia} & c_i + \sum_a v_a T_{ia} < c_m \end{cases} \tag{S9}$$

#### 1.3 Tumor evolution and phenotype equations

Tumor evolution is governed by a mutation rate, which can randomly shift a cell phenotype up or down by one increment along two independent phenotypic axes: Angiogenesis phenotype $a$ and glucose consumption phenotype $g$. Each phenotype can shift between 5 states, which we will represent as coordinate pairs ($a$,$g$). At state (0,0), the phenotype rates are equivalent to that of epithelial cells. Each increment of the glucose consumption phenotype changes the glucose consumption rate of glucose $r_g$, by a power of 2 (Equation S10) and each increment of the angiogenesis phenotype changes the angiogenesis rate $v_a$ by a power of 2, with a jump from zero angiogenesis at the first state (Equation S11). At the first increment, this rate is set so that a voxel completely filled with tumor cells would increase its endothelial carrying capacity by one endothelial carrying-capacity unit, defined as the baseline endothelial carrying capacity of maximally dense normal tissue.

$$r_g = r_e \, 2^g \tag{S10}$$

$$v_a = \begin{cases} 0 & a = 0 \\ v_1 \, 2^{a-1} & a > 0 \end{cases} \tag{S11}$$

### 2 Additional patient data analysis

The 45 case dataset used in this study is generated from 30 individual patients, and 32 tumors, with two pairs of tumors derived from single patients. 13 of the tumors were tracked for 3 subsequent years, meaning that two cases could be drawn from each of these. We looked briefly at how the parameter fits changed over years in Figure S4 A-D, and how the percent volumes change over years in Figure S4E. Though there appears to be some connection between first and second year results, we focused our analysis on the 45 cases independently to increase our sample size and study the model's ability to represent one year of tumor growth. We also looked for differences in growth rate and central density between squamous and adenocarcinoma in our dataset (Figure S4F,G).

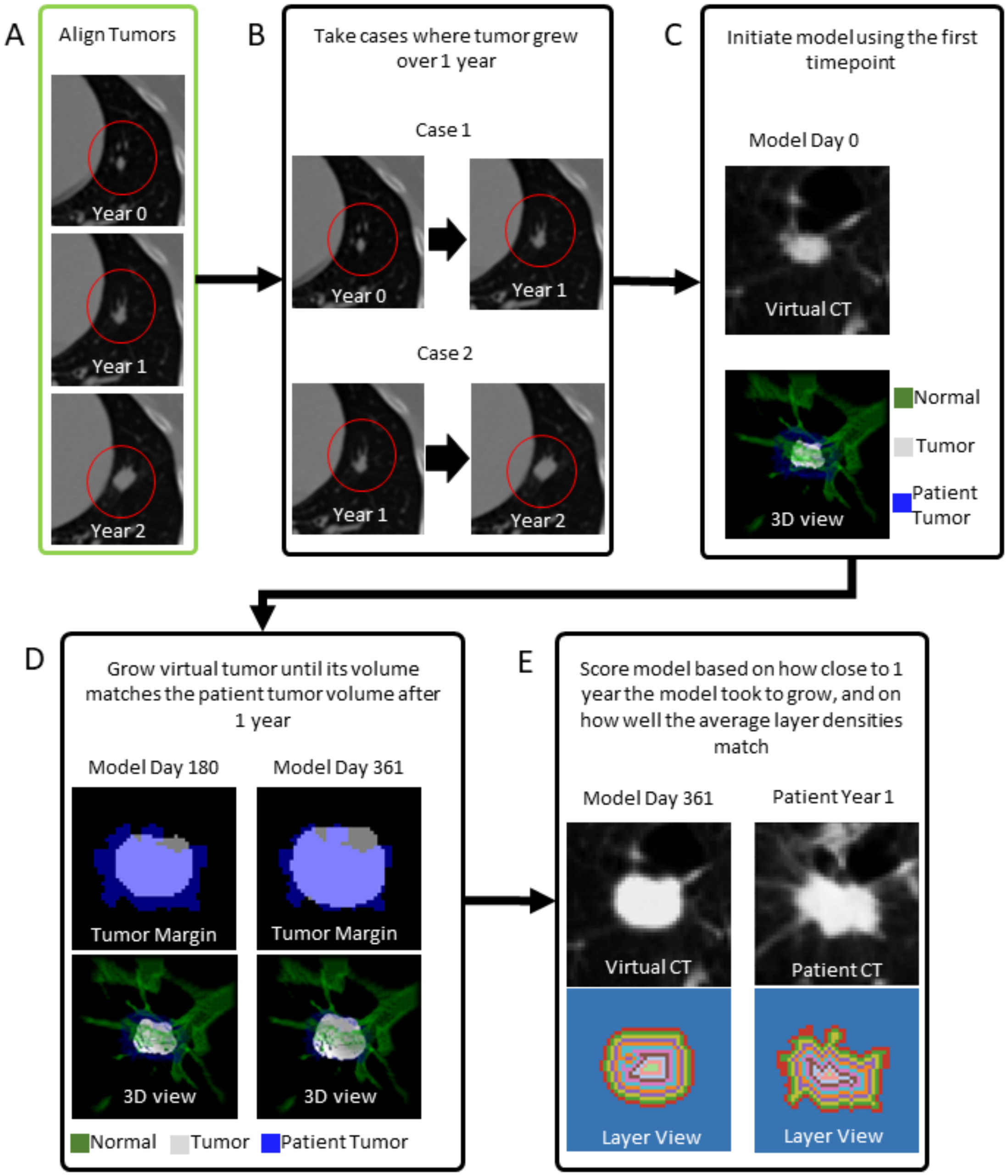


Supplementary Figure 3: **A** The patient data was first processed by finding series of consecutive year measurements and ensuring that the same tumor was segmented in each. **B** Aligned tumors that grew over 1 year became cases that the model would be scored against. **C** The initial model timepoint. At this stage the model looks identical to the CT scan. **D** Shows how the tumor volume changes over time and approaches the volume of the patient tumor after 1 year. **E** The model is scored based two criteria: on how close to 1 year of simulated time it took the model to achieve the same volume as the patient tumor after 1 year; on how similar the average density of each matching layer of the model and patient tumors are. Colors in the Layer View indicate matching layers.

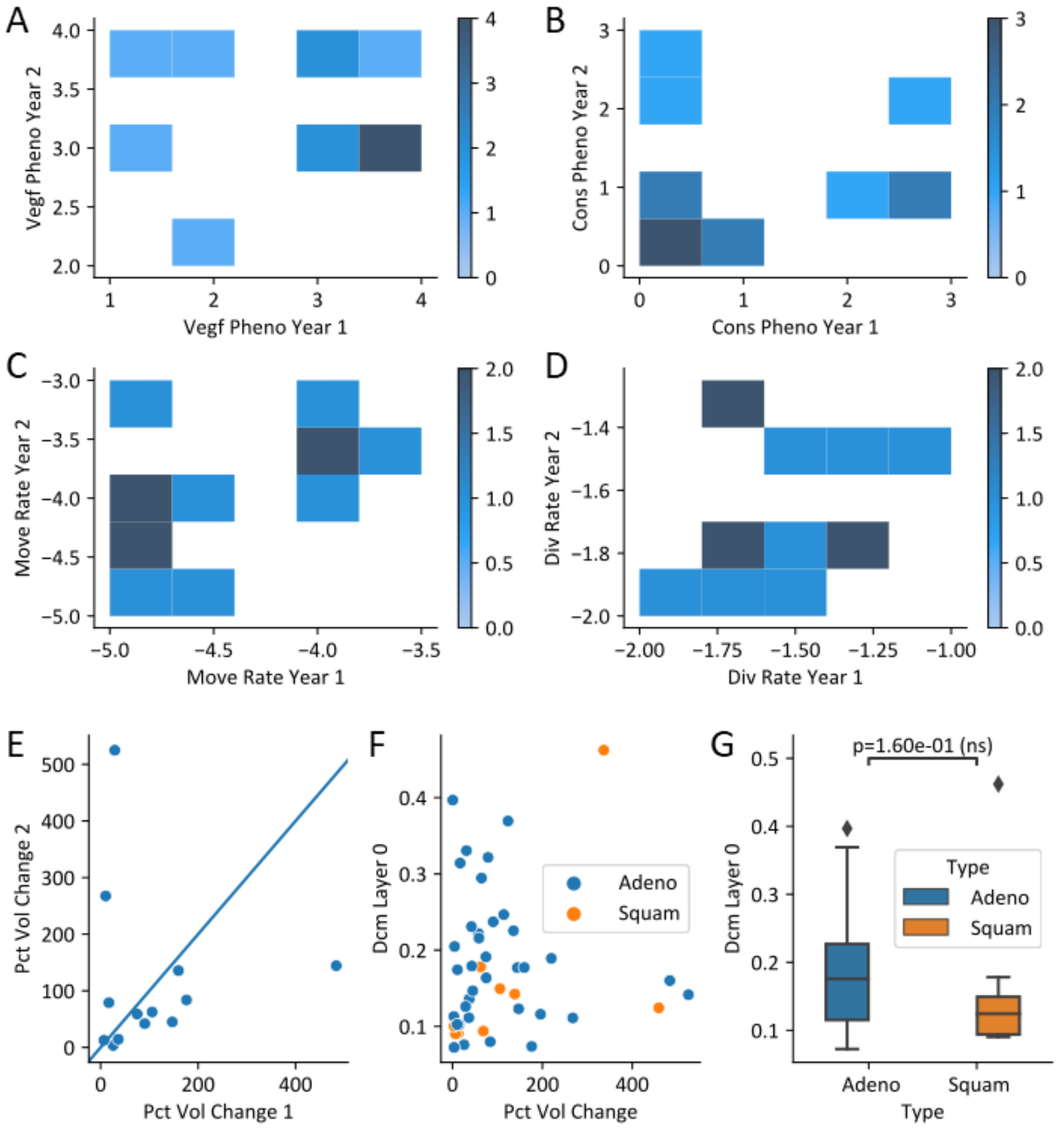


Supplementary Figure 4: **A-D** Best fitting phenotypes for the 13 patients whose tumor was tracked for 2 years. **E** Percent volume change between year 1 and year 2. **F** Differences in density of the central tumor layer and the percent volume change of the tumor from all 45 cases is shown to compare squamous and adenocarcinoma phenotypes. **G** Box and whisker representation of the density of the central tumor layer to look for statistical significance.
